# SRPK1 is a determinant of chemoresistance to Docetaxel in prostate cancer

**DOI:** 10.64898/2026.09.16.751805

**Authors:** Duygu Duzgun, William Read, Sebastian Oltean

**Author notes:** Correspondence: Prof Sebastian Oltean, Department of Clinical and Biomedical Sciences, University of Exeter Medical School, St Luke’s Campus, Exeter, EX12LU, UK.

## Abstract

Docetaxel is a major therapeutic option for advanced prostate cancer, but the development of acquired resistance substantially limits its clinical efficacy. Although multiple mechanisms have been implicated in docetaxel resistance, the upstream regulatory pathways coordinating these phenotypes remain incompletely understood. Here, we investigated the role of serine/arginine-rich protein kinase 1 (SRPK1), a key regulator of pre-mRNA splicing, in acquired docetaxel resistance in prostate cancer. Docetaxel-resistant PC3 cells showed increased SRPK1 expression at both the protein and RNA levels. Genetic depletion or pharmacological inhibition of SRPK1 substantially restored docetaxel sensitivity, whereas ectopic SRPK1 expression in parental PC3 cells increased resistance. Mechanistically, SRPK1 inhibition reduced expression of βIII-tubulin and restored docetaxel-induced microtubule bundling. SRPK1 inhibition also increased apoptosis in resistant cells, accompanied by increased cleavage of caspase-8, caspase-9 and PARP. In parallel, SRPK1 inhibition restored E-cadherin expression and reduced the enhanced migratory phenotype of resistant cells. At the signalling and RNA-processing levels, docetaxel-resistant cells exhibited increased EGFR expression and increased phosphorylation of the SRPK1 substrate SRSF1. SRSF1 depletion partially restored docetaxel sensitivity, supporting an EGFR-SRPK1-SRSF1 pathway in resistance. Importantly, inhibition of SRPK1 or depletion of SRSF1 altered the splicing of apoptosis– and microtubule-associated genes, increasing the pro-apoptotic Bcl-xS and MCL-1S isoforms and shifting tau splicing towards 3R at the expense of 4R. Collectively, these findings identify SRPK1 as a central regulator of multiple phenotypic and molecular features of docetaxel resistance in prostate cancer. Targeting SRPK1-dependent splicing may therefore represent a strategy to re-sensitize resistant prostate cancer cells to docetaxel.

## 1. Introduction

Prostate cancer is one of the most frequently diagnosed malignancies in men and remains a major cause of cancer-related mortality. Although localized disease can often be effectively managed, progression to advanced or metastatic disease remains a major therapeutic challenge. In particular, metastatic castration-resistant prostate cancer (mCRPC) remains incurable and is characterized by the emergence of resistance to multiple therapeutic approaches. Docetaxel, a taxane that stabilizes microtubules and disrupts mitotic progression, has been a key component of systemic therapy for advanced prostate cancer and provides a significant survival benefit in patients with mCRPC. However, primary and acquired resistance to docetaxel substantially limits its effectiveness [1–3].

Docetaxel binds to β-tubulin within microtubules and promotes microtubule polymerization while suppressing their normal dynamic instability. This results in abnormal spindle formation, mitotic arrest and, ultimately, cell death. Consequently, alterations in microtubule composition and dynamics are important mechanisms of taxane resistance. In prostate cancer, increased expression of βIII-tubulin has been associated with reduced sensitivity to docetaxel and with adverse clinical outcomes, providing evidence that changes in the microtubule machinery can contribute directly to therapeutic resistance [4–6]. Additional mechanisms include increased drug efflux, alterations in androgen receptor signalling, PI3K/AKT activation, changes in apoptotic signalling and acquisition of stem-like or mesenchymal phenotypes [2,3].

Epithelial–mesenchymal transition (EMT) has also been implicated in docetaxel resistance. Acquisition of mesenchymal characteristics can enhance migration, invasion, survival and resistance to apoptosis. In prostate cancer models, docetaxel-resistant cells have been reported to exhibit reduced E-cadherin and increased expression of mesenchymal and stem-like features [7]. Thus, the development of chemoresistance is likely to involve coordinated changes in several cellular processes rather than a single resistance mechanism.

An additional layer of regulation that has received comparatively little attention in chemotherapy resistance is alternative pre-mRNA splicing. Alternative splicing enables a single gene to generate multiple mRNA and protein isoforms and is extensively deregulated in cancer. Serine/arginine-rich protein kinase 1 (SRPK1) is a central regulator of this process. SRPK1 phosphorylates serine/arginine-rich splicing factors, including SRSF1, thereby regulating their activity, localization and splice-site selection [8,9]. SRPK1 expression or activity is increased in several malignancies and has been linked to oncogenic signalling, tumour progression and altered alternative splicing [8–10].

Our previous work established that SRPK1 is increased in prostate cancer and regulates alternative splicing of VEGF-A, promoting expression of pro-angiogenic isoforms [11,12]. These observations raised the possibility that SRPK1 may also regulate alternative splicing programmes involved in cancer cell survival and therapeutic resistance. In support of this concept, inhibition of SRPK1 has been shown in other cancer models to alter the splicing of apoptosis-associated genes, including BCL2L1 and MCL1, resulting in increased production of pro-apoptotic isoforms [13].

Here, we investigated the role of SRPK1 in acquired docetaxel resistance in prostate cancer. Using parental and docetaxel-resistant PC3 cells, together with genetic manipulation and pharmacological inhibition of SRPK1, we demonstrate that SRPK1 is a functional determinant of docetaxel resistance. We further show that SRPK1 regulates multiple resistance-associated phenotypes, including βIII-tubulin expression and microtubule responses, apoptosis and EMT-associated migration. Finally, our data identify an EGFR–SRPK1–SRSF1 signalling axis and demonstrate that SRPK1-dependent splicing of apoptosis– and microtubule-associated transcripts provides a mechanistic link between altered RNA processing and docetaxel resistance.

## 2. Materials and Methods

### 2.1 Cell culture and generation of docetaxel-resistant cells

Human prostate cancer PC3 and LNCaP cell lines were used in this study. Parental cells were maintained under standard tissue-culture conditions in the appropriate complete culture medium supplemented with fetal bovine serum and antibiotics.

Docetaxel-resistant derivatives were generated by progressive exposure of parental prostate cancer cells to increasing concentrations of docetaxel, with surviving cells maintained and expanded during selection. Resistant cells were subsequently maintained under drug-free conditions for experimental analysis unless otherwise indicated. The resistant PC3 population was designated PC3-DtxR, while parental cells were designated PC3-P.

### 2.2 Docetaxel sensitivity and cell viability assays

Parental and docetaxel-resistant cells were seeded at equal density and exposed to increasing concentrations of docetaxel. Cell viability was determined using an MTT-based colorimetric assay following the experimental protocol used in the study.

Dose-response curves were generated and IC50 values were calculated by nonlinear regression. Where indicated, cells were treated with docetaxel in combination with the SRPK1 inhibitor SPHINX31 or following genetic depletion of SRPK1 or SRSF1.

### 2.3 SRPK1 inhibition and gene knockdown

SRPK1 activity was inhibited pharmacologically using SPHINX31. Appropriate vehicle-treated controls were included in all experiments. For genetic depletion, cells were transfected with SRPK1-specific small interfering RNA (siRNA) together with an appropriate non-targeting control siRNA.

SRSF1 expression was similarly depleted using SRSF1-specific siRNA. Knockdown efficiency was assessed by western blotting and/or RT-PCR as appropriate.

### 2.4 SRPK1 overexpression

Parental PC3 cells were transfected with an expression plasmid encoding SRPK1 or the corresponding control plasmid. Cells were selected/expanded as appropriate and SRPK1 expression was confirmed by western blotting. The effect of SRPK1 overexpression on docetaxel sensitivity was subsequently determined by MTT assay.

### 2.5 RNA isolation and RT-PCR

Total RNA was extracted from parental and resistant prostate cancer cells using a standard RNA isolation procedure (Trizol). RNA was reverse transcribed to generate cDNA, followed by PCR amplification of the transcripts of interest.

SRPK1 transcript abundance was assessed to determine differences between parental and resistant cells. Alternative splicing of BCL2L1, MCL1 and MAPT was assessed using isoform-specific or splice-sensitive RT-PCR assays. Products corresponding to alternative splice isoforms were resolved by electrophoresis and quantified where appropriate.

### 2.6 Western blotting

Cells were lysed in appropriate protein extraction buffer containing protease and phosphatase inhibitors. Protein concentration was determined using a standard protein assay. Equal quantities of protein were resolved by SDS-PAGE and transferred to membranes. GelDoc easy (BioRad) was used to assess and normalize equal loading.

Membranes were incubated with antibodies against SRPK1, βIII-tubulin, E-cadherin, EGFR, SRSF1 and phosphorylated SRSF1, together with antibodies recognizing cleaved caspase-8, cleaved caspase-9 and cleaved PARP where indicated. Appropriate loading controls were used. Following incubation with HRP-conjugated secondary antibodies, proteins were detected by enhanced chemiluminescence.

Band intensities were quantified using appropriate densitometric analysis and normalized to the relevant loading control.

### 2.7 Immunofluorescence analysis of microtubules

Parental and docetaxel-resistant cells were treated with docetaxel with or without SPHINX31. Cells were fixed, permeabilized and incubated with an antibody against βIII-tubulin, followed by an appropriate fluorescent secondary antibody.

Microtubule organization was examined by fluorescence microscopy. In parental cells, docetaxel treatment produced the characteristic concentrated/bundled microtubule staining pattern. Resistant cells displayed a predominantly diffuse cytoplasmic βIII-tubulin pattern despite docetaxel exposure, whereas combined SRPK1 inhibition and docetaxel treatment restored the microtubule-bundling phenotype.

### 2.8 Measurement of apoptosis

Changes in mitochondrial membrane potential, as an early indicator of apoptosis, were assessed using the JC-10 assay. Parental and resistant cells were treated with increasing concentrations of docetaxel in the presence or absence of SRPK1 inhibition.

Apoptosis was further assessed by western blotting for cleavage of caspase-8, caspase-9 and PARP.

### 2.9 Migration assay

Cell migration was assessed using a wound-healing/scratch assay. Confluent monolayers of parental, resistant and SRPK1-manipulated cells were wounded and imaged at defined time points following treatment.

Migration was quantified by measuring the change in wound area over time. Where indicated, resistant cells were treated with SPHINX31 or subjected to SRPK1 knockdown.

### 2.10 Statistical analysis

Data are presented as mean ± SEM or mean ± SD, as appropriate. Statistical analyses were performed using Prism. Comparisons between two groups were performed using an appropriate two-tailed statistical test, while experiments involving multiple groups were analysed using one-way or two-way ANOVA with an appropriate post hoc test.

For dose-response experiments, IC50 values were calculated by nonlinear regression. A P value <0.05 was considered statistically significant.

## 3. Results

### 3.1 SRPK1 is overexpressed in Docetaxel-resistant prostate cancer cells and is involved in the resistance mechanism

To be able to study the role of SRPK1 in chemoresistance in prostate cancer (PCa), PC3 cell lines resistant to Docetaxel (one of the most common treatments in PCa) have been obtained (see Methods). The resistant PC3 cell line (named PC3-DtxR) has an IC50 to Docetaxel almost 20 times higher than the parental PC3 (**Figure 1A**). Western blot and RT-PCR analysis showed that SRPK1 is overexpressed at both protein and RNA level in the resistant cells compared to the parental ones (**Figure 1B and Supplementary Figure 1**).

**Figure 1.**
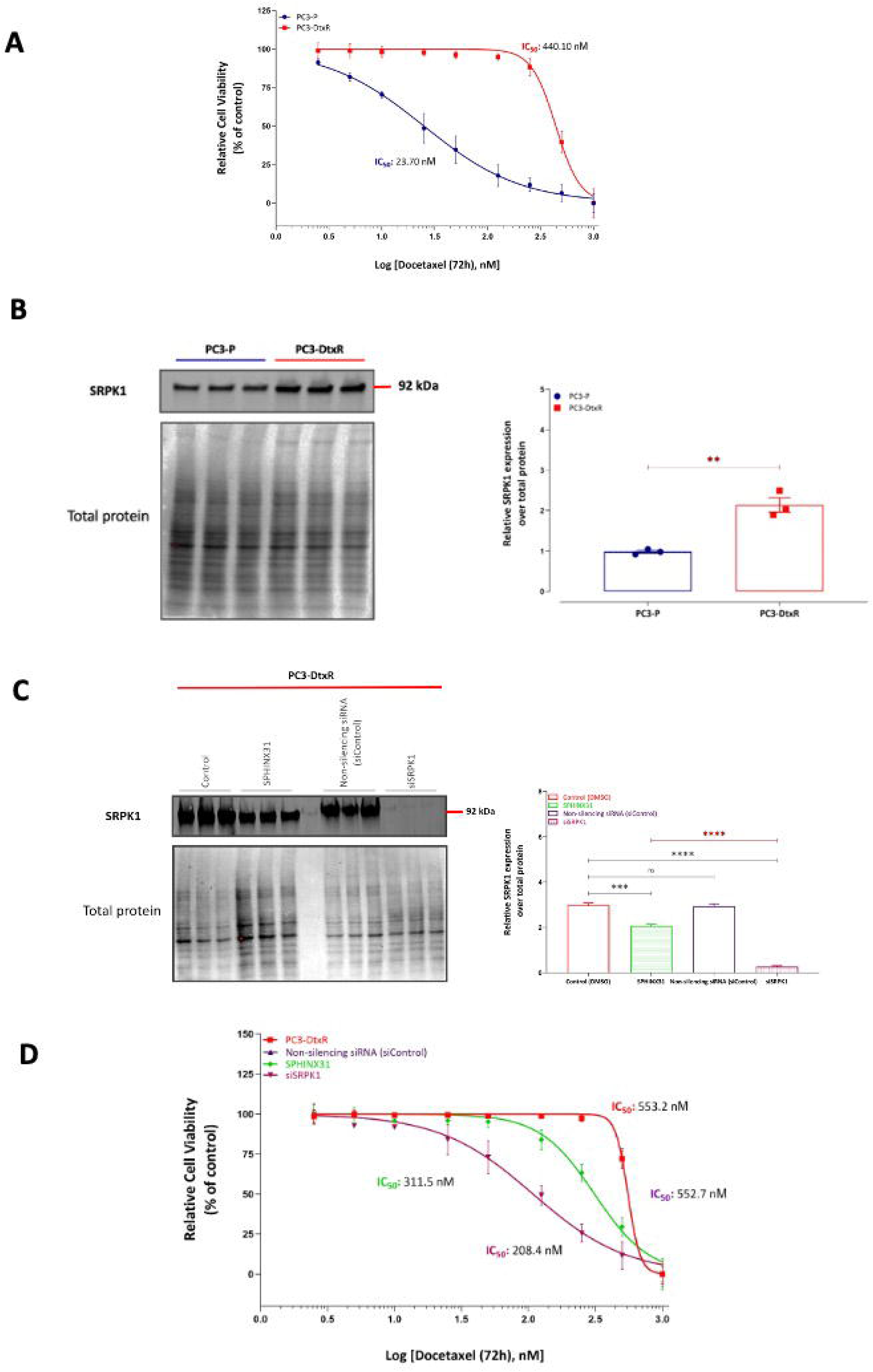
SRPK1 is overexpressed in Docetaxel-resistant PC3 cells and its chemical or genetic inhibition rescues the resistant phenotyope. (A) Docetaxel IC50 curves obtained with an MTT assay for sensitive (parental) PC3 cell (PC3-P) and resistant ones (PC3-DtxR) (B) Western blot analysis shows that SRPK1 is increased in resistant cells. Left upper blot – SRPK1 antibody; Left lower blot – total protein quantified on the transfer membrane using BioRad Geldoc; right graph – western blot quantification (C) Western blot and quantification showing a robust knockdown of SRPK1 in PC3 resistant cells (D) IC50 for Docetaxel is significantly decreased when SRPK1 is inhibited either genetically (knockdown) or chemically (SPHINX31)

To test whether SRPK1 is involved in the mechanism of resistance to Docetaxel or a byproduct of the cells transformation, we knocked down SRPK1 expression in resistant cells as well as treated them with SRPK1 inhibitor SPHINX31 (**Figure 1C**). As shown in **Figure 1D**, both SRPK1 knock-down as well as chemical inhibition of SRPK1 results in a pronounced rescue of the IC50, making the cells more sensitive to Docetaxel.

All the above evidence suggests that SRPK1 is involved mechanistically in the development of Docetaxel chemoresistance in PC3 cells.

### 3.2 Overexpression of SRPK1 in parental (sensitive) PC3 cells confers resistance to Docetaxel comparable to the PC3-resistant cells

To explore whether exogenous overexpression of SRPK1 in sensitive PC3 cells induces resistance to Docetaxel we transfected PC3 sensitive (parental) cells with an SRPK1 plasmid. **Figure 2, left panel** shows Western blot analysis for three PC3-parental cell lines (non-transfected, transfected with control plasmid, transfected with SRPK1 plasmid) and the resistant PC3. Exogenous expression of SRPK1 from the plasmid increased SRPK1 level by about two-fold compared to the parental controls and is about half of the expression level in the resistant cell line. In **Figure 2, right panel**, the cell viability was evaluated by MTT assay. Exogenous overexpression of SRPK1 increases IC50 for Docetaxel to about half the value of resistant cells, in concordance with the expression levels. This suggests that SRPK1 is a strong determinant of resistance to Docetaxel in PC3 cells.

**Figure 2.**
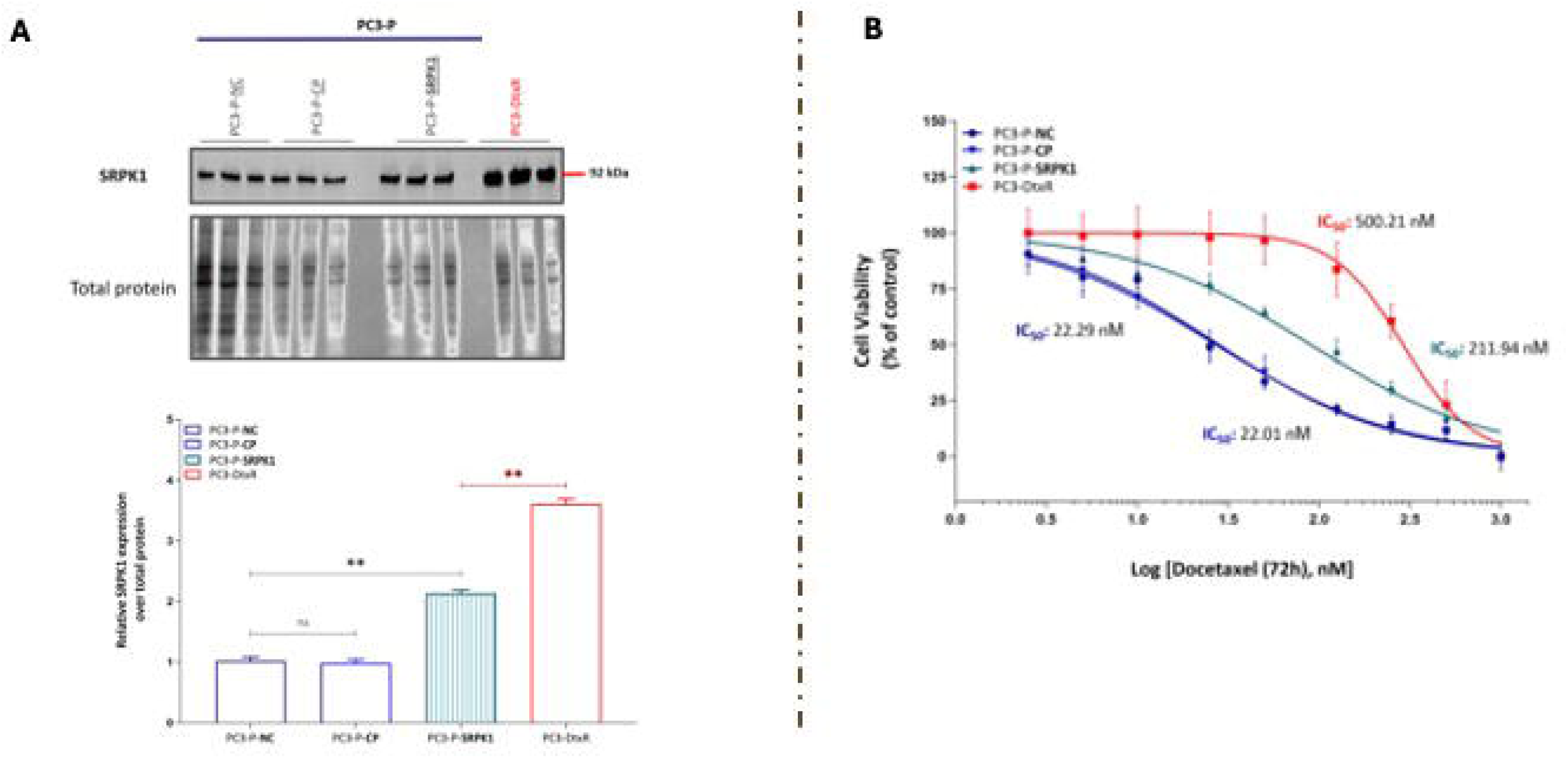
Exogenous overexpression of SRPK1 in PC3 parental (sensitive) cells confers resistance to Docetaxel. **(A)** Western blot for SRPK1 showing expression levels across several cell lines: PC3-parental (non-transfected, transfected with control plasmid, transfected with SRPK1 plasmid) and the resistant PC3. Left lower blot – total protein quantified on the transfer membrane using BioRad Geldoc; graph below – western blot quantification **(B)** IC50 determinations in PC3 parental cell line, PC3 parental with exogenous SRPK1 overexpression and PC3 resistant cell line

**Figure 3.**
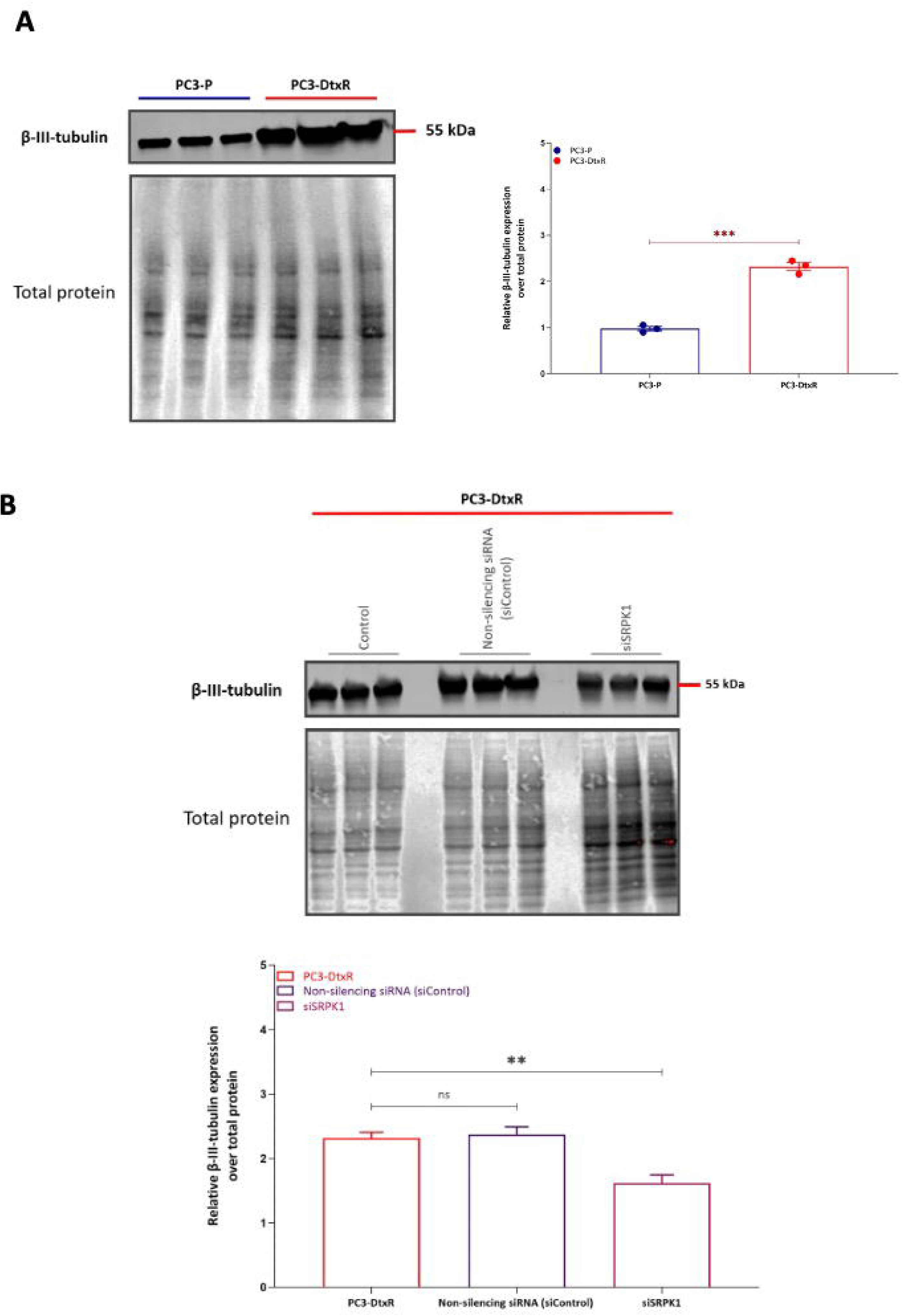
Silencing of SRPK1 in Docetaxel-resistant PC3 cells reduces expression of βIII-tubulin. **(A)** Western blot for βIII-tubulin expression in sensitive (parental) PC3 cell (PC3-P) and resistant ones (PC3-DtxR); Left lower blot – total protein quantified on the transfer membrane using BioRad Geldoc; right graph – western blot quantification **(B)** Western blot for βIII-tubulin expression in resistant PC3 cells – from left, control, transfected with non-silencing siRNA and transfected with SRPK1-silencing siRNA; panel below the blot – total protein quantified on the transfer membrane using BioRad Geldoc; right graph – western blot quantification

### 3.3 SRPK1 modulates the Docetaxel microtubules resistance mechanism

We further wanted to explore whether SRPK1 modulates known Docetaxel resistance mechanisms. One of these mechanisms involves the microtubules – Docetaxel promotes and stabilizes microtubules by binding to the beta-tubulin subunits of microtubules, promoting their assembly and preventing their depolymerization. This disrupts the natural dynamic equilibrium between tubulin and microtubules inside the cell. Changes in the composition or structure of microtubules, such as the upregulation of βIII-tubulin, can make them less sensitive to Docetaxel’s stabilizing effects.

Indeed, when probed for expression, βIII-tubulin was increased in resistant cells compared to sensitive ones (**Figure 2A**) and knockdown of SRPK1 in resistant cells results in decreased expression of βIII-tubulin (Figure **2B**).

Staining for microtubules with βIII-tubulin antibodies shows a diffuse cytoplasmic pattern when microtubules are intact. When cells are exposed to Docetaxel, there is “microtubule bundling” in the cytoplasm, meaning that immunofluorescence staining for βIII-tubulin shows a focused and punctuate pattern.

Indeed, as shown in **Supplementary Figure 2**, βIII-tubulin staining shows microtubule bundling when exposed to Docetaxel, however, the staining is diffuse cytoplasmic in resistant cells, even when exposed to Docetaxel. When an inhibitor of SRPK1 is added to the Docetaxel treatment, microtubule bundling is seen again, suggesting that SRPK1 is involved in this resistance mechanism.

### 3.4 SRPK1 modulates apoptosis in Docetaxel-resistant cells

Another mechanism of resistance to Docetaxel involves modulation of apoptosis in cancer cells. To investigate whether SRPK1 interferes with this mechanism, a JC 10 assay (which measures mitochondrial membrane potential and therefore evaluates initial stages of apoptosis) was employed. As shown in **Figure 4A, left panel**, increasing doses of Docetaxel increase apoptosis in PC3 parental cells (PC3-P); when SRPK1 is exogenously overexpressed in PC3-P cells, there is leass increase in apoptosis at the same concentrations of Docetaxel – see **Figure 4A, right panel**.

**Figure 4.**
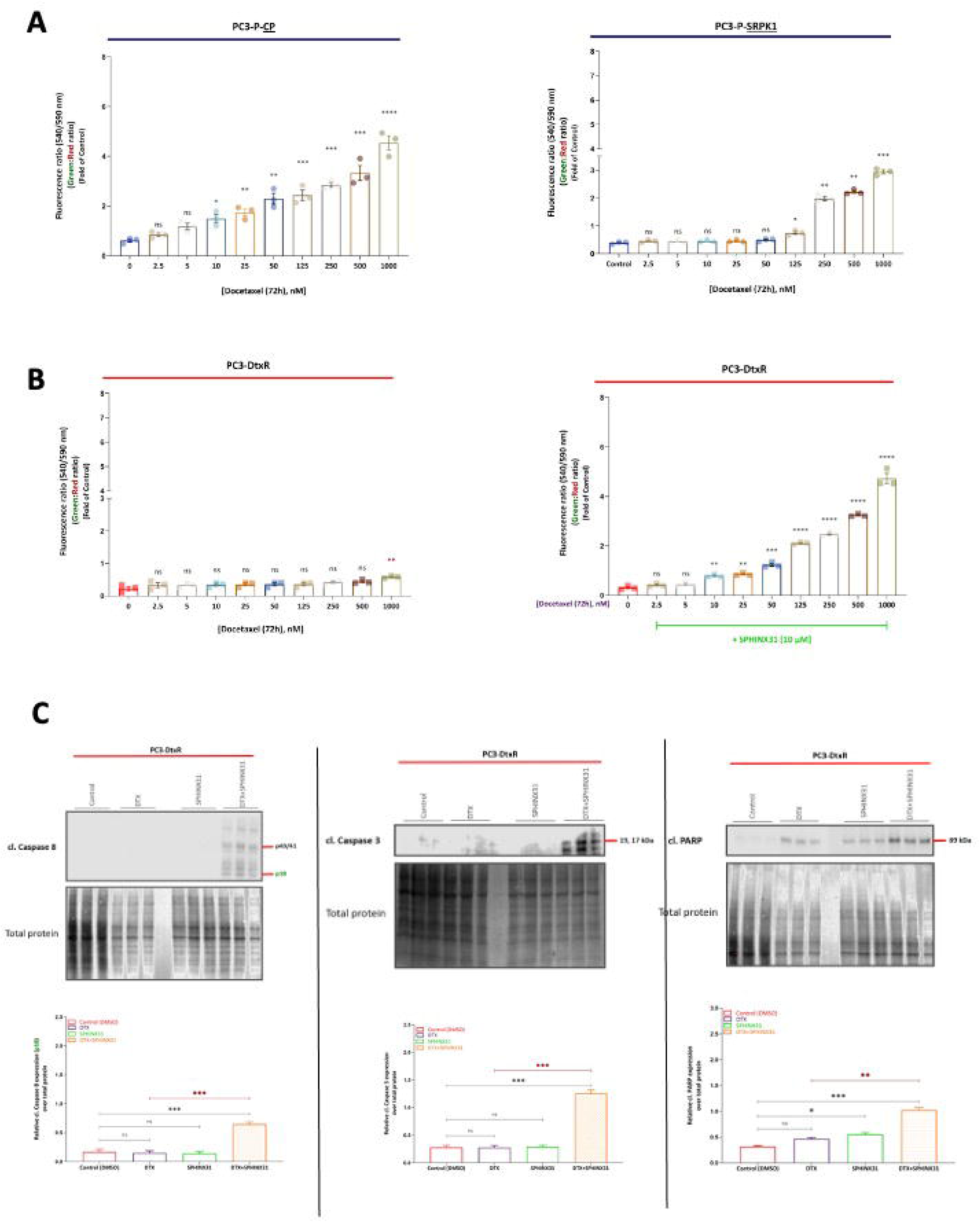
Pharmacological inhibition of SRPK1 with SPHINX31 re-sensitises PC3-DtxR cells to docetaxel and promotes apoptosis. **(A)** Dose-dependent changes in fluorescence ratio following treatment with increasing concentrations of docetaxel (DTX; 72 h) in parental PC3-P cells, PC3-P cells overexpressing SRPK1 (PC3-P-SRPK1), and **(B)** PC3-DtxR cells in the absence or presence of SPHINX31 (10 μM). Fluorescence ratios are expressed relative to the untreated control. **(C)** Western blot analysis of apoptotic markers in PC3-DtxR cells treated with DTX, SPHINX31, or the combination of DTX and SPHINX31. Cleaved caspase-8, cleaved caspase-3 and cleaved PARP were assessed, with total protein used as a loading control. Quantification of the corresponding protein expression relative to total protein is shown below each immunoblot. Data are presented as mean ± SEM. Statistical significance is indicated by asterisks; ns, not significant.

Similar doses of Docetaxel do not induce apoptosis even at high values in PC3-resistant cells – **Figure 4B, left panel**, while co-treatment with the SRPK1 inhibitor Sphinx clearly sensitizes these cells to induce apoptosis – **Figure 4B, right panel**.

We also evaluated the cleaved products of caspase 8, 9 and PARP; as shown in **Figure 4C**, addition of the SRPK1 inhibitor Sphinx 31 to Docetaxel treatment clearly increases apoptosis (appearance of cleaved products – see last lane 4 in each panel) suggesting that this re-sensitizes the PC3-resistant cells.

### 3.5 SRPK1 modulates epithelial-mesenchymal transitions and migration in Docetaxel-resistant cells

Yet another mechanism described for Docetaxel resistance is modulation of epithelial-mesenchymal transitions (EMT), with resistant cells moving towards a more mesenchymal / stem-like phenotype.

To test involvement of SRPK1 in this mechanism we first assessed expression of E-cadherin, a junctional protein in epithelial cells whose expression is decreased when cells transition to mesenchymal phenotypes and it is the most commonly used marker for EMT. As seen in **Figure 5A**, there is a marked decrease in E-cadherin expression in resistant cells in comparison to the parental ones; inhibition of SRPK1 with SPHINX31 on its own or together with Docetaxel restores strong expression of E-cadherin – see **Figure 5B**.

**Figure 5.**
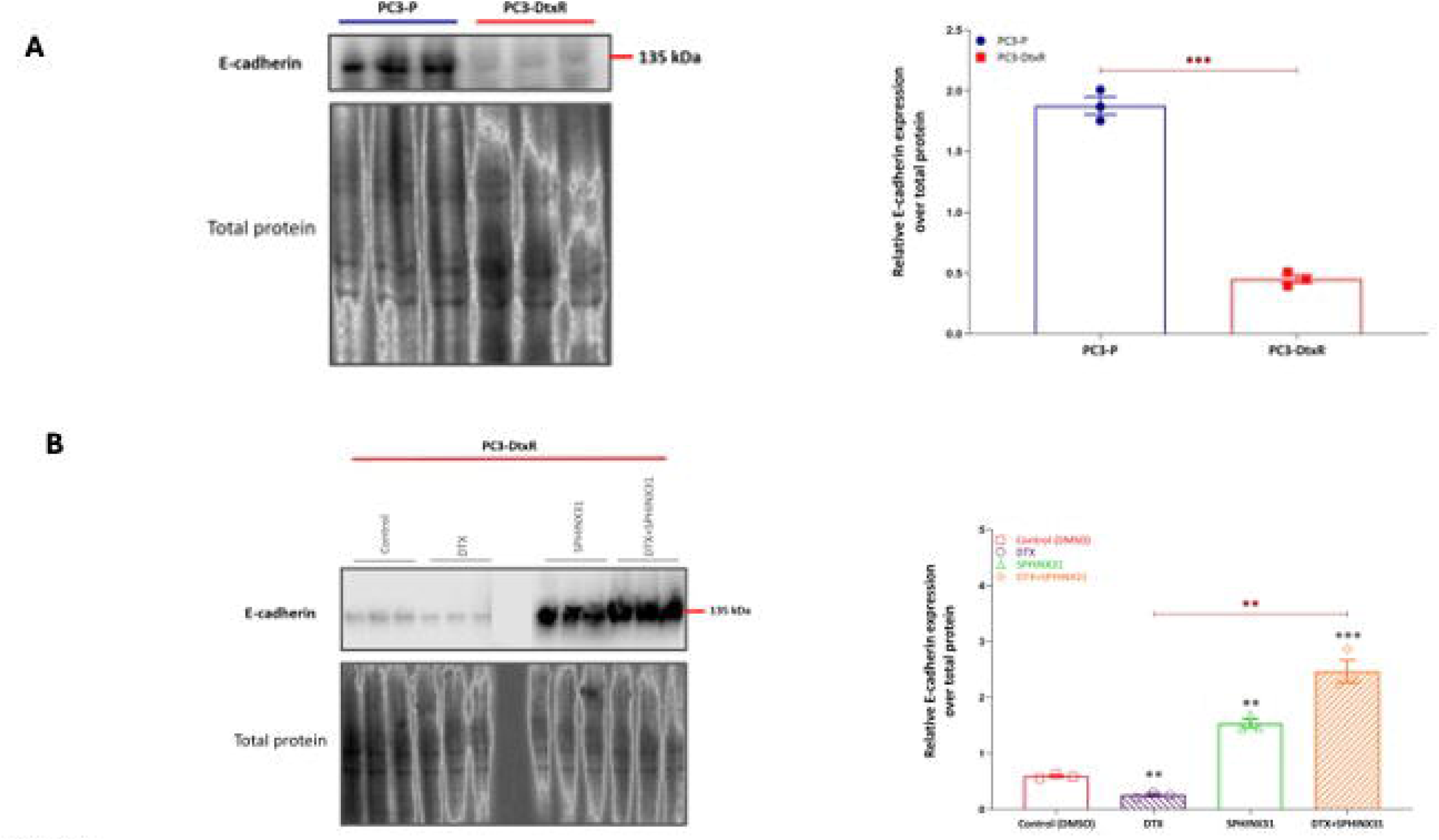
SPHINX31 restores E-cadherin expression in docetaxel-resistant PC3-DtxR cells. **(A)** Western blot analysis of E-cadherin expression in parental PC3-P and docetaxel-resistant PC3-DtxR cells. Quantification of E-cadherin expression relative to total protein is shown on the right. **(B)** Western blot analysis of E-cadherin expression in PC3-DtxR cells following treatment with DTX, SPHINX31, or the combination of DTX and SPHINX31. Quantification of E-cadherin expression relative to total protein is shown on the right. Data are presented as mean ± SEM. Statistical significance is indicated by asterisks; ns, not significant.

If SRPK1 affects EMT the functional expectation is that there will be a modulation of migration in resistant cells. Indeed, as seen in **Figure 6A**, Docetaxel-resistant cells as well as parental cell with overexpression of SRPK1 have increased migration; either chemical (**Figure 6B**) or genetic (**Supplementary Figure 3**) inhibition of SRPK1 decreases migration of resistant cells.

**Figure 6.**
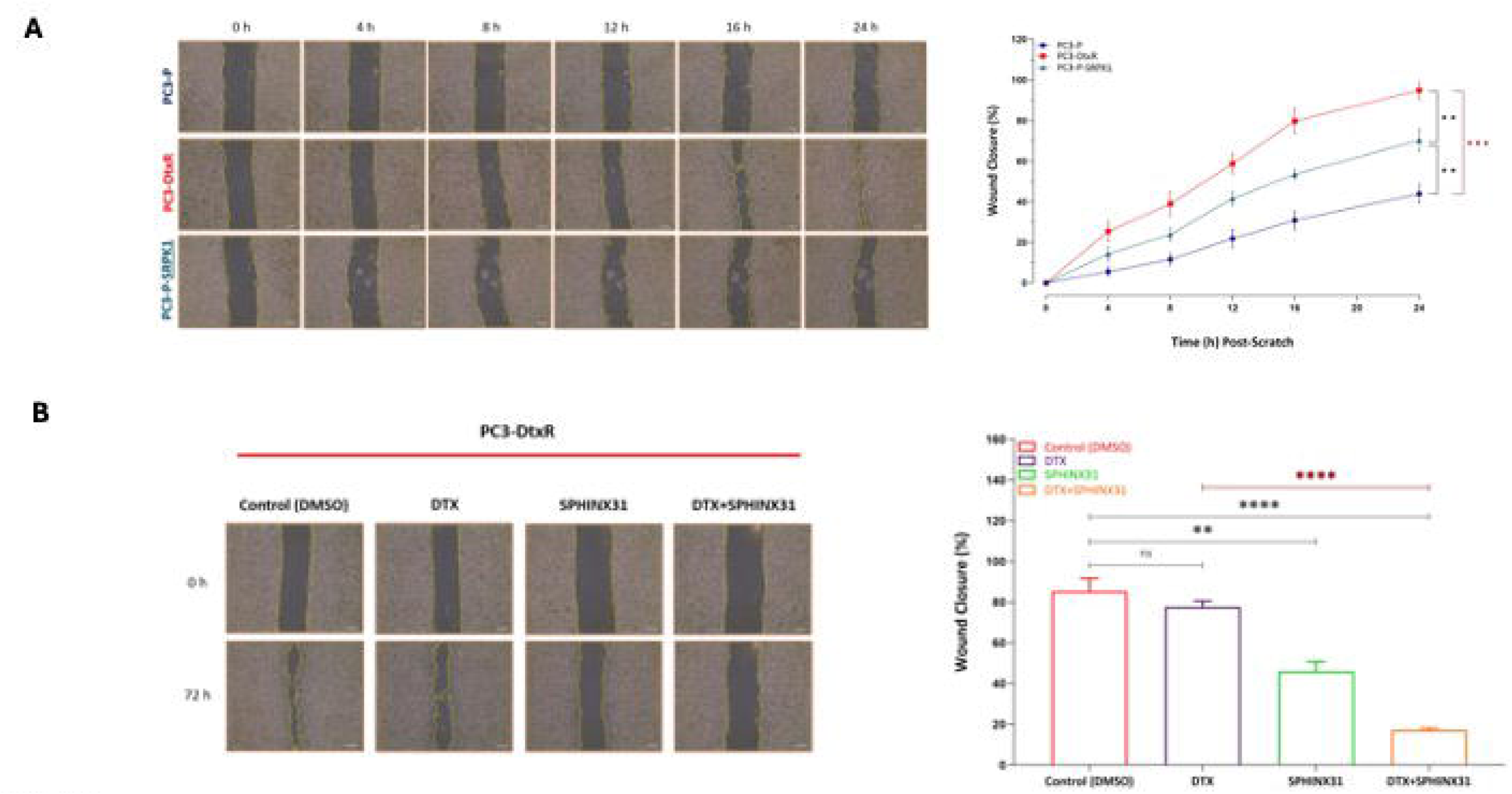
SRPK1 expression enhances migration of prostate cancer cells and SPHINX31 inhibits migration of PC3-DtxR cells. (**A**) Representative images from scratch-wound assays performed in parental PC3-P, docetaxel-resistant PC3-DtxR and PC3-P-SRPK1 cells at 0, 4, 8, 12, 16 and 24 h following scratch formation. Quantification of wound closure over time is shown on the right. (B) Representative images of scratch wounds in PC3-DtxR cells treated with vehicle control (DMSO), DTX, SPHINX31, or the combination of DTX and SPHINX31 at 0 and 72 h. Quantification of wound closure at 72 h is shown on the right. Data are presented as mean ± SEM. Statistical significance is indicated by asterisks; ns, not significant.

### 3.6 SRPK1-related resistance mechanism involves EGFR and splicing factor SRSF1

Following establishment of the involvement of SRPK1 in the resistance to Docetaxel, we further wanted to explore what signalling pathways and which splice factors may be involved. SRPK1 has been reported before to be activated in the EGFR/ Akt pathway and one of its most common substrate that it phosphorylates and activates is SRSF1. We therefore tested the involvement of EGFR and SRSF1.

As seen in **Figure 7A** – EGFR is upregulated in Docetaxel-resistant cells; SRSF1 is hyper-phosphorylated (**Figure 7B**) and knockdown of SRSF1 (**Figure 7C** and **Supplementary Figure 4**) results in partial rescue of the Docetaxel resistance (see MTT assay in **Figure 7D**).

**Figure 7.**
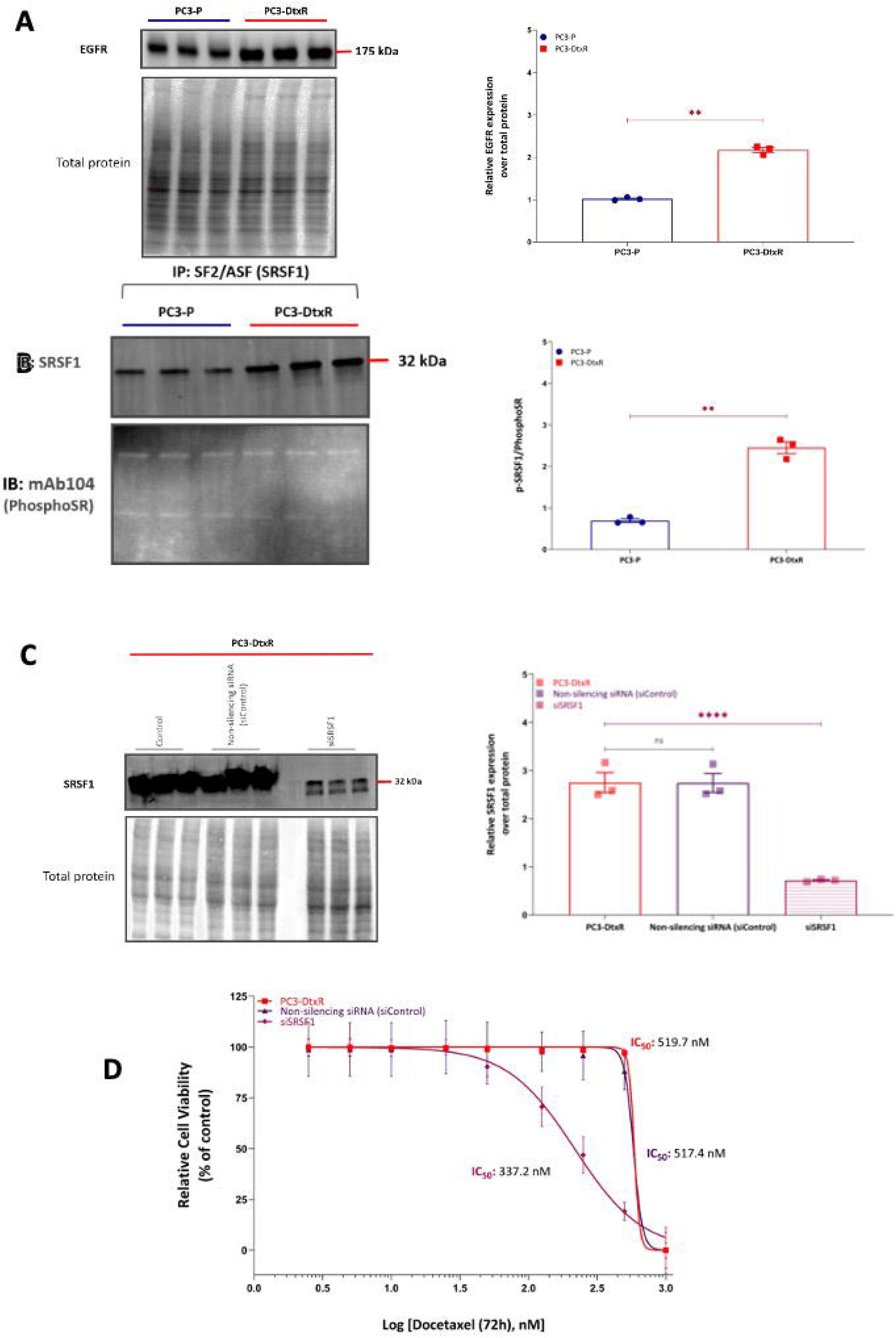
SRPK1-dependent phosphorylation of SRSF1 contributes to docetaxel resistance in PC3-DtxR cells. (**A**) Western blot analysis of EGFR expression in parental PC3-P and docetaxel-resistant PC3-DtxR cells, with quantification of EGFR expression relative to total protein shown on the right. (**B**) Immunoprecipitation of SRSF1 from PC3-P and PC3-DtxR cells followed by immunoblotting with an anti-phosphoserine antibody (mAb104) to assess SRSF1 phosphorylation. SRSF1 immunoblotting was used to assess immunoprecipitation efficiency. Quantification of SRSF1 phosphorylation relative to SRSF1 expression is shown on the right. (**C**) Western blot analysis showing SRSF1 expression following transfection of PC3-DtxR cells with non-silencing control siRNA or siRNA targeting SRSF1. Quantification relative to total protein is shown on the right. (**D**) Dose-response curves showing the effect of increasing concentrations of docetaxel on cell viability in PC3-DtxR cells, non-silencing siRNA-treated cells and SRSF1-knockdown cells. The corresponding IC₅₀ values are indicated. Data are presented as mean ± SEM. Statistical significance is indicated by asterisks; ns, not significant.

All of this evidence suggests that the axis EGFR-SRPK1-SRSF1 is activated in Docetaxel-resistant cells and forms a major mechanism of the resistance.

### 3.7 SRPK1 and SRSF1 control Docetaxel resistance through changing splicing patterns of key genes

SRPK1, as a splicing kinase, and SRSF1, as a splice factor, control splicing patterns of many genes. We reasoned that their effect on various mechanisms of Docetaxel resistance must be through changing specific splice isoforms ratios.

Indeed, as seen in **Figure 8A and B**, in the presence of Docetqxel and Sphinx31, proapoptotic splice isoforms Bcl-Xs and Mcl15-s increase expression; this will trigger and increase in apoptosis and cell death therefore the sensitization effect. Tau 3R and 4R are two major splice isoforms of the microtubule-associated protein tau, defined by having either three (3R) or four (4R) carboxy-terminal binding repeat domains. 4R tau binds to microtubules more strongly and has a higher capacity to stabilize them compared to 3R tau. As seen in **Figure 8C**, when Docetaxel is combined with Sphinx31 there is an increase in 3R and decrease in 4R, indicating microtubule de-stabilization and therefore re-sensitizationj to Docetaxel.

**Figure 8.**
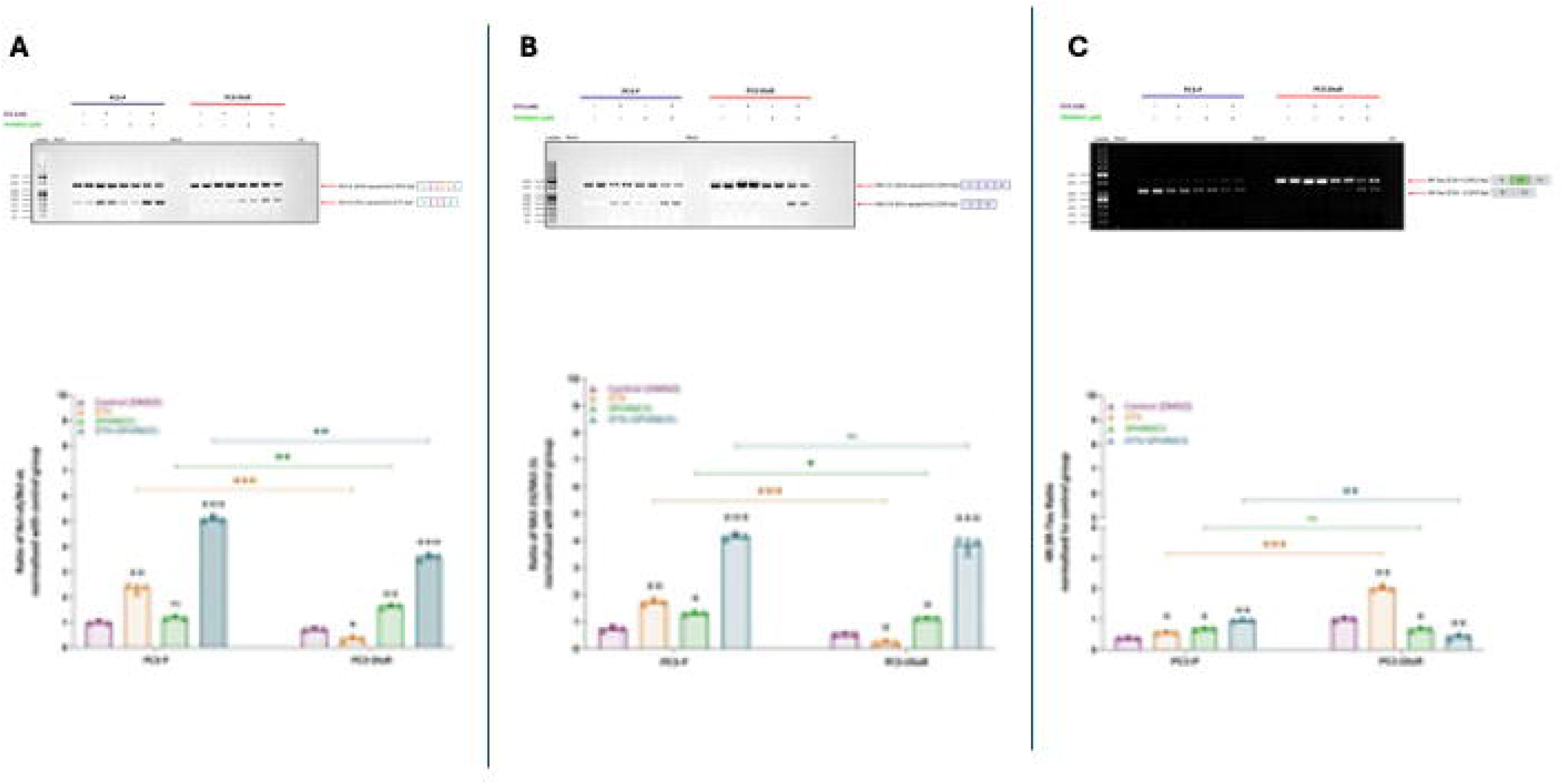
SPHINX31 modulates alternative splicing of apoptosis– and survival-associated transcripts in PC3-P and PC3-DtxR cells. (**A**) Representative RT-PCR analysis of alternative splicing of Bcl-x, showing the pro-survival Bcl-xL and pro-apoptotic Bcl-xS isoforms in PC3-P and PC3-DtxR cells following treatment with vehicle control (DMSO), DTX, SPHINX31, or the combination of DTX and SPHINX31. Quantification of the Bcl-xS/Bcl-xL ratio, normalised to the control group, is shown below. (**B**) Representative RT-PCR analysis of Mcl-1 alternative splicing, showing the anti-apoptotic Mcl-1L and pro-apoptotic Mcl-1S isoforms. Quantification of the Mcl-1S/Mcl-1L ratio is shown below. (**C**) Representative RT-PCR analysis of Tau alternative splicing showing the 4R-Tau and 3R-Tau isoforms. Quantification of the 4R-Tau/3R-Tau ratio, normalised to the control group, is shown below. Data are presented as mean ± SEM. Statistical significance is indicated by asterisks; ns, not significant.

Similar to inhibition of SRPK1 activity, the same effects are seen when knocking down SRSF1 (See **Supplementary Figures 5, 6 and 7**).

## 4. Discussion

The development of resistance to docetaxel represents a major obstacle to the effective treatment of advanced prostate cancer. Although several mechanisms of resistance have been identified, including alterations in tubulin, activation of survival pathways, drug efflux and acquisition of mesenchymal or stem-like characteristics, the mechanisms linking these phenotypes remain incompletely understood [1–3]. Our findings identify SRPK1 as a central regulator of docetaxel resistance and suggest that altered RNA splicing provides an important mechanistic connection between several otherwise distinct resistance phenotypes.

The first important observation was the marked increase in SRPK1 expression in docetaxel-resistant PC3 cells. Resistant cells exhibited an approximately 20-fold increase in the docetaxel IC50 compared with parental cells, accompanied by increased SRPK1 expression at both the RNA and protein levels. Importantly, this association was supported by functional experiments. Depletion of SRPK1 or pharmacological inhibition with SPHINX31 substantially restored docetaxel sensitivity, whereas ectopic expression of SRPK1 in parental cells increased the IC50. Thus, the increased expression of SRPK1 in resistant cells is unlikely to represent merely a consequence of selection or adaptation to prolonged drug exposure; rather, SRPK1 appears to contribute directly to the resistant phenotype.

This finding extends previous observations linking SRPK1 to prostate cancer biology. SRPK1 is increased in prostate cancer tissue and has previously been shown to regulate VEGF-A alternative splicing and tumour angiogenesis [11,12]. The current findings suggest that the biological importance of SRPK1 in prostate cancer extends beyond angiogenesis and includes regulation of tumour-cell responses to cytotoxic therapy. More broadly, SRPK1 has emerged as an important regulator of cancer-associated splicing programmes and has been implicated in several oncogenic signalling pathways [8–10].

One mechanism through which SRPK1 appears to regulate docetaxel resistance is the microtubule response. Docetaxel acts primarily through stabilization of microtubules, and alterations in tubulin composition can reduce its effectiveness. In particular, βIII-tubulin has been repeatedly associated with taxane resistance, including in prostate cancer [4–6]. In our model, βIII-tubulin was increased in docetaxel-resistant cells, while SRPK1 depletion reduced βIII-tubulin expression. Furthermore, resistant cells failed to display the characteristic microtubule-bundling response to docetaxel, whereas inhibition of SRPK1 restored this phenotype. These observations place SRPK1 upstream of a clinically relevant mechanism of taxane resistance.

The ability of SRPK1 to influence microtubule behaviour may also involve alternative splicing of microtubule-associated proteins. Our data show that inhibition of SRPK1 shifts tau expression from the 4R towards the 3R isoform. Tau isoforms differ in their effects on microtubule dynamics, with 4R tau having a greater capacity to stabilize microtubules than 3R tau [15,16]. Although the precise contribution of tau isoform switching to docetaxel resistance requires further investigation, the observed shift is consistent with a model in which SRPK1-dependent splicing promotes a microtubule state that is more permissive to resistance, whereas SRPK1 inhibition restores a more docetaxel-responsive state.

A second major phenotype regulated by SRPK1 was apoptosis. Docetaxel treatment increased apoptosis in parental PC3 cells, whereas resistant cells were comparatively refractory to apoptotic induction. SRPK1 overexpression in parental cells reduced the apoptotic response to docetaxel, while inhibition of SRPK1 restored apoptosis in resistant cells. This was supported by increased cleavage of caspase-8, caspase-9 and PARP following combined docetaxel and SPHINX31 treatment. These results indicate that SRPK1 contributes to resistance not only by modifying the cellular target of docetaxel but also by suppressing the downstream execution of cell death.

The alternative splicing results provide a potential mechanistic explanation for this effect. SRPK1 inhibition increased the relative abundance of the pro-apoptotic Bcl-xS and MCL-1S isoforms. BCL2L1 and MCL1 are well-established examples of genes in which alternative splicing determines the balance between pro– and anti-apoptotic protein isoforms [13,14]. Bcl-xL and MCL-1L favour cell survival, whereas Bcl-xS and MCL-1S promote apoptosis. The observation that both pharmacological inhibition of SRPK1 and depletion of SRSF1 produced similar changes in these splice patterns strongly supports a model in which SRPK1 regulates resistance through an SRSF1-dependent splicing programme.

SRSF1 is a well-established substrate of SRPK1 and a major regulator of alternative splicing. SRPK1-mediated phosphorylation controls SRSF1 localization and splicing activity [8,9]. In our resistant cells, EGFR was increased and SRSF1 displayed increased phosphorylation. Importantly, depletion of SRSF1 partially restored docetaxel sensitivity. Together, these findings support an EGFR–SRPK1–SRSF1 signalling axis in docetaxel-resistant prostate cancer.

The relationship between EGFR signalling and SRPK1 is particularly interesting because activation of EGFR and downstream PI3K/AKT signalling has previously been implicated in prostate cancer therapeutic resistance. SRPK1 can function downstream of oncogenic signalling pathways, and AKT has been reported to regulate SRPK1 activation [10,17]. Thus, EGFR activation may provide an upstream signal that enhances SRPK1/SRSF1 activity, resulting in coordinated changes in alternative splicing that favour survival during docetaxel exposure. Further experiments will be required to establish the precise hierarchy of EGFR, AKT, SRPK1 and SRSF1 in the resistant cells.

SRPK1 also regulated the mesenchymal phenotype. Docetaxel-resistant cells displayed reduced E-cadherin expression and increased migration, consistent with the association between EMT and taxane resistance described in prostate cancer [7]. Inhibition or depletion of SRPK1 restored E-cadherin expression and reduced migration. Moreover, ectopic SRPK1 expression in parental cells increased migration. These observations suggest that SRPK1 contributes to the broader aggressive phenotype associated with treatment-resistant prostate cancer. Whether SRPK1 directly controls EMT through alternative splicing of specific EMT regulators or indirectly through changes in survival signalling remains to be determined.

Taken together, the results support a model (see **Figure 9**) in which SRPK1 sits at the intersection of several major components of docetaxel resistance. Increased EGFR signalling is associated with increased SRPK1 activity and SRSF1 phosphorylation. The resulting alteration in alternative splicing favours anti-apoptotic isoforms such as Bcl-xL and MCL-1L and a microtubule-associated tau isoform profile enriched for 4R tau. In parallel, SRPK1 activity is associated with increased βIII-tubulin expression, reduced docetaxel-induced microtubule bundling, suppression of apoptosis and acquisition of a more migratory/mesenchymal phenotype. Pharmacological or genetic inhibition of SRPK1 reverses several of these changes, thereby restoring sensitivity to docetaxel.

**Figure 9.**
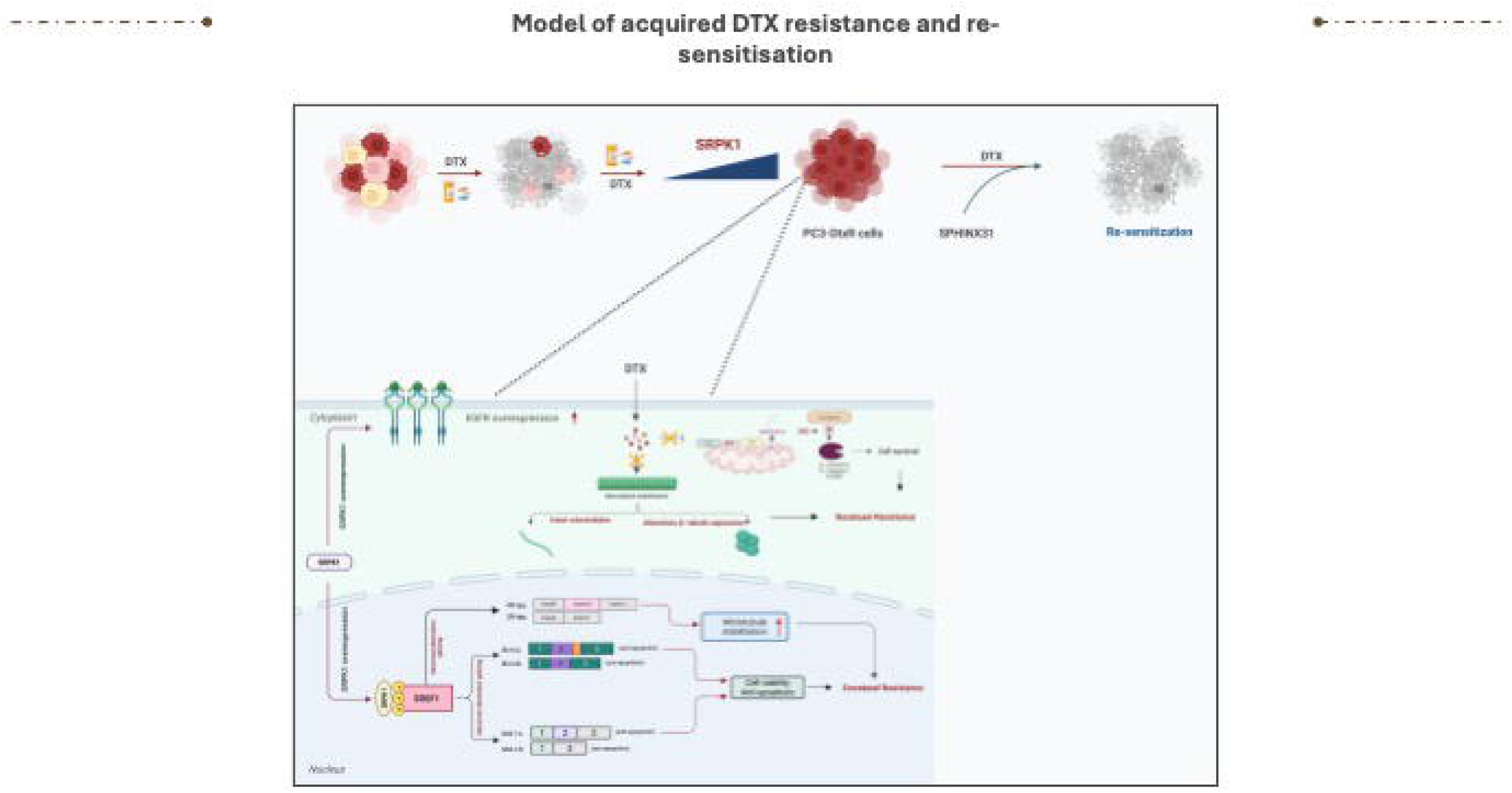
Proposed model of acquired docetaxel resistance and re-sensitisation. Schematic model summarising the proposed mechanisms underlying acquired docetaxel (DTX) resistance in PC3-DtxR cells and their re-sensitisation following SRPK1 inhibition. Chronic DTX exposure is associated with increased SRPK1 expression in cancer cells with elevated EGFR expression, which results in enhanced SRSF1 phosphorylation. Increased SRPK1/SRSF1 activity promotes aberrant alternative splicing of transcripts involved in microtubule stability and apoptosis, including Tau, Bcl-x and Mcl-1, thereby favouring microtubule stabilisation, cell survival and resistance to DTX. Pharmacological inhibition of SRPK1 with SPHINX31 reverses these changes, enhances apoptotic responses and restores DTX sensitivity. The model also incorporates changes in β-tubulin expression and microtubule stability associated with the resistant phenotype.

An important implication of these findings is the potential therapeutic value of combining SRPK1 inhibition with docetaxel. The current data show that SRPK1 inhibition alone can reverse several resistance-associated phenotypes and, importantly, markedly sensitizes resistant cells to docetaxel. This provides a rationale for investigating SRPK1 inhibitors as combination therapies in docetaxel-resistant prostate cancer. SPHINX31 and related SRPK-targeting compounds have already been used experimentally to modulate SR-protein phosphorylation and alternative splicing in cancer models [13,14]. However, further work will be required to establish therapeutic windows, specificity and efficacy in vivo.

Several limitations should also be considered. First, the central functional experiments presented here are primarily based on the PC3 model, and validation in additional prostate cancer models, will strengthen the generality of the findings. Second, although the results strongly support an SRPK1/SRSF1-dependent mechanism, direct demonstration that individual splice isoforms are necessary and sufficient for resistance will require isoform-specific rescue or overexpression experiments. Third, the relationship between EGFR activation and SRPK1 activity requires further mechanistic dissection, particularly to determine whether EGFR/AKT signalling directly controls SRPK1 activation in the resistant cells. Finally, validation of SRPK1 expression, SRSF1 phosphorylation and the relevant splice isoform changes in clinical samples from patients before and after docetaxel treatment will be important to determine whether this pathway has predictive or therapeutic relevance in patients.

In conclusion, our findings identify SRPK1 as a functional determinant of docetaxel resistance in prostate cancer. SRPK1 inhibition restores docetaxel sensitivity by simultaneously affecting microtubule responses, apoptosis and EMT-associated phenotypes, with SRSF1-dependent alternative splicing providing a plausible molecular mechanism linking these processes. The EGFR–SRPK1–SRSF1 axis therefore represents a potentially targetable pathway for overcoming docetaxel resistance and warrants further investigation as a therapeutic strategy in advanced prostate cancer.

## Acknowledgements

This research was funded by a PhD studentship to Duygu Duzgun awarded by the Ministry of National Education, The Republic of Turkey and by BBSRC grant BB/ J007293 / 2 to SO.

