## Supplementary files for "SRPK1 is a determinant of chemoresistance to Docetaxel in prostate cancer"

### Slide 1
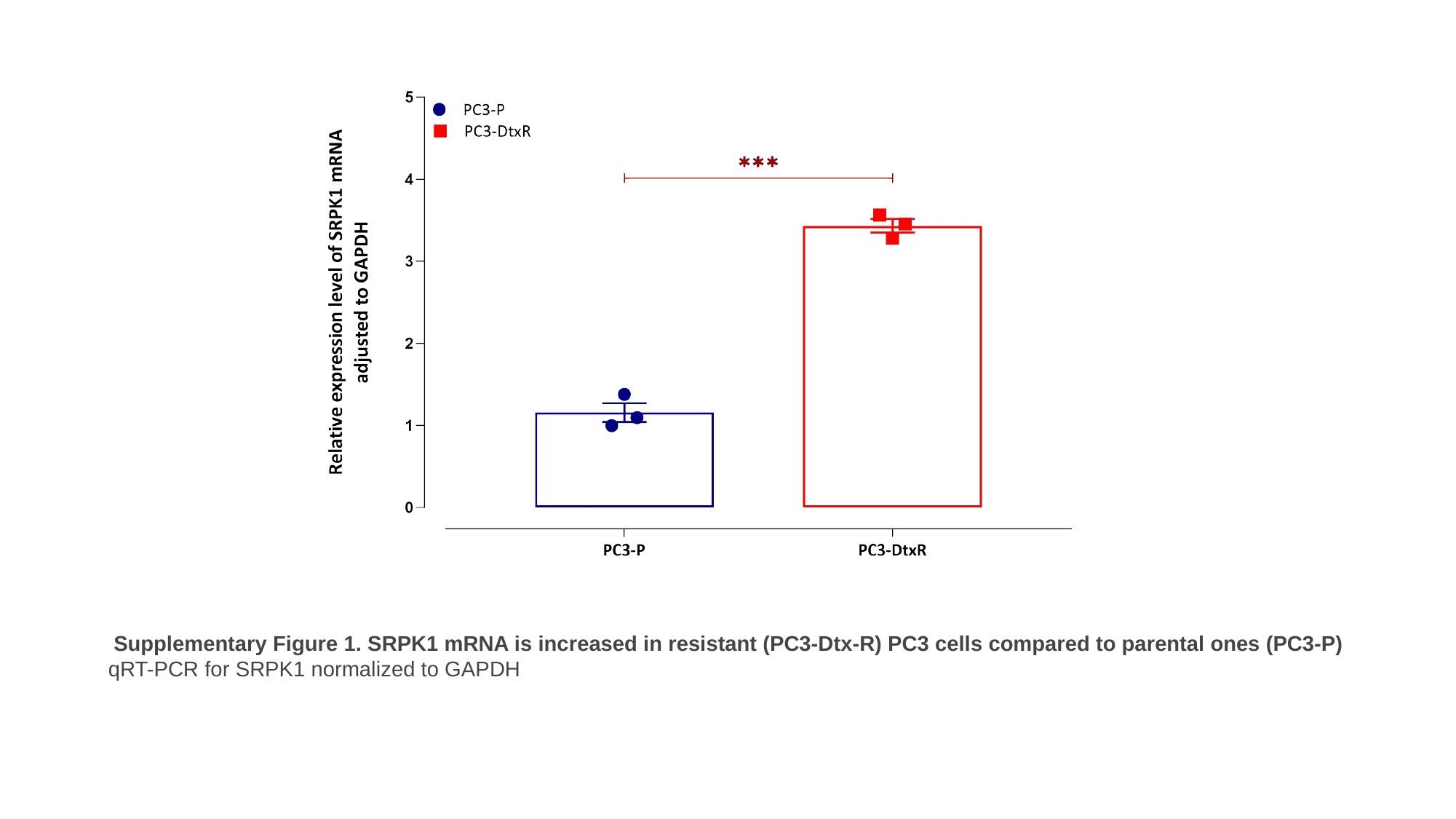

Supplementary Figure 1. SRPK1 mRNA is increased in resistant (PC3-Dtx-R) PC3 cells compared to parental ones (PC3-P)
qRT-PCR for SRPK1 normalized to GAPDH

### Slide 2
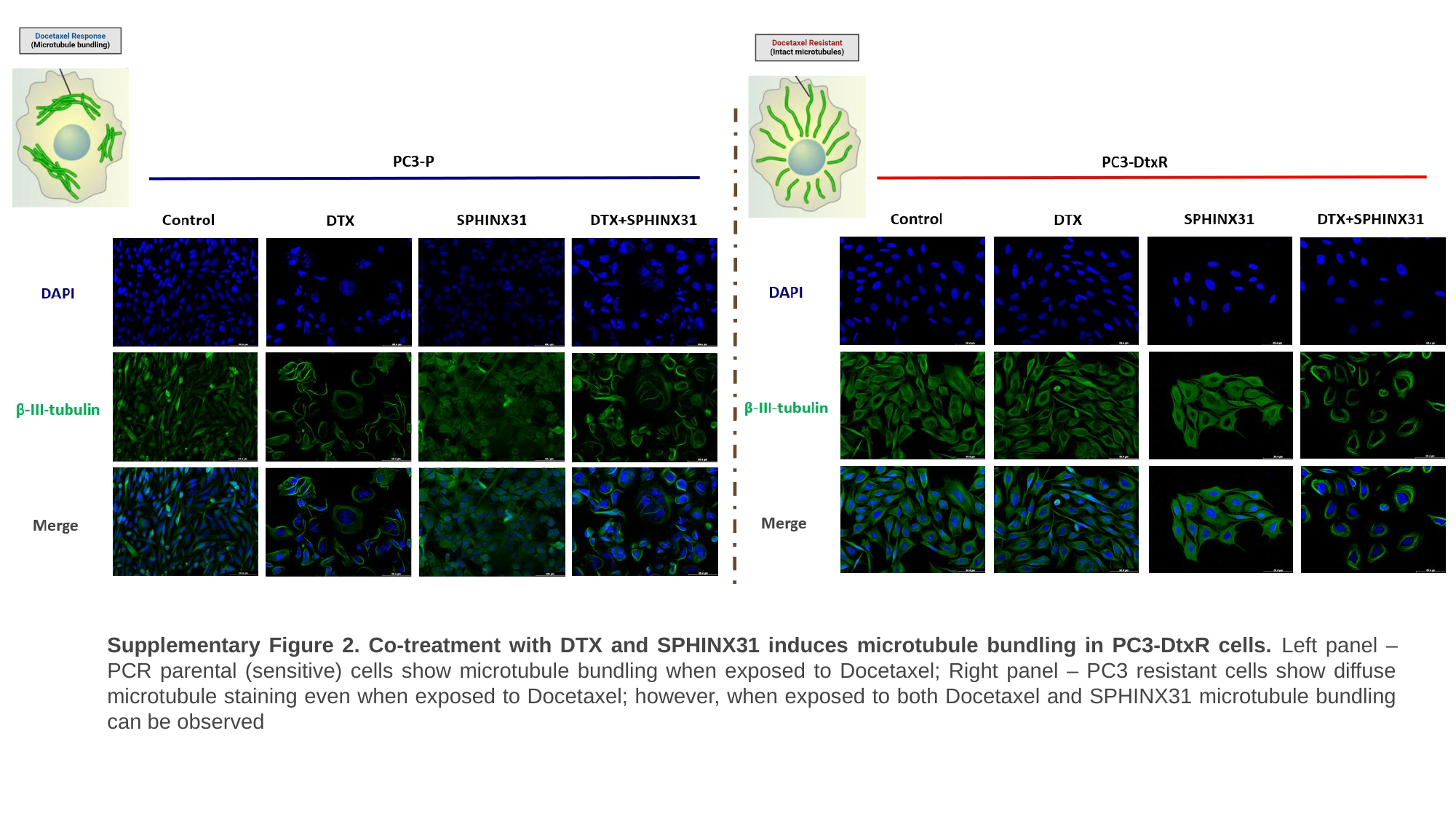

Supplementary Figure 2. Co-treatment with DTX and SPHINX31 induces microtubule bundling in PC3-DtxR cells. Left panel – PCR parental (sensitive) cells show microtubule bundling when exposed to Docetaxel; Right panel – PC3 resistant cells show diffuse microtubule staining even when exposed to Docetaxel; however, when exposed to both Docetaxel and SPHINX31 microtubule bundling can be observed

### Slide 3
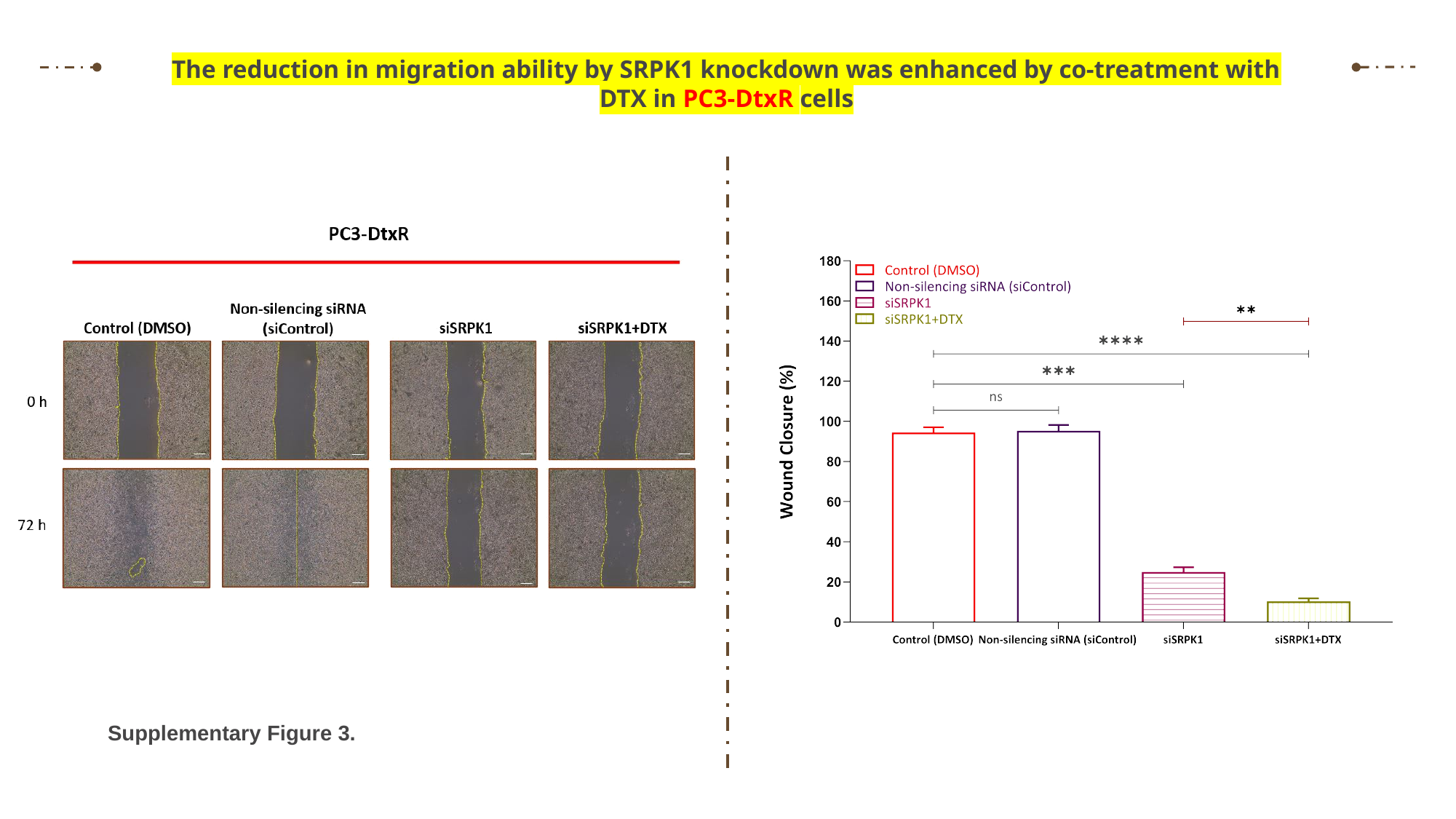

The reduction in migration ability by SRPK1 knockdown was enhanced by co-treatment with DTX in PC3-DtxR cells
Supplementary Figure 3.

### Slide 4
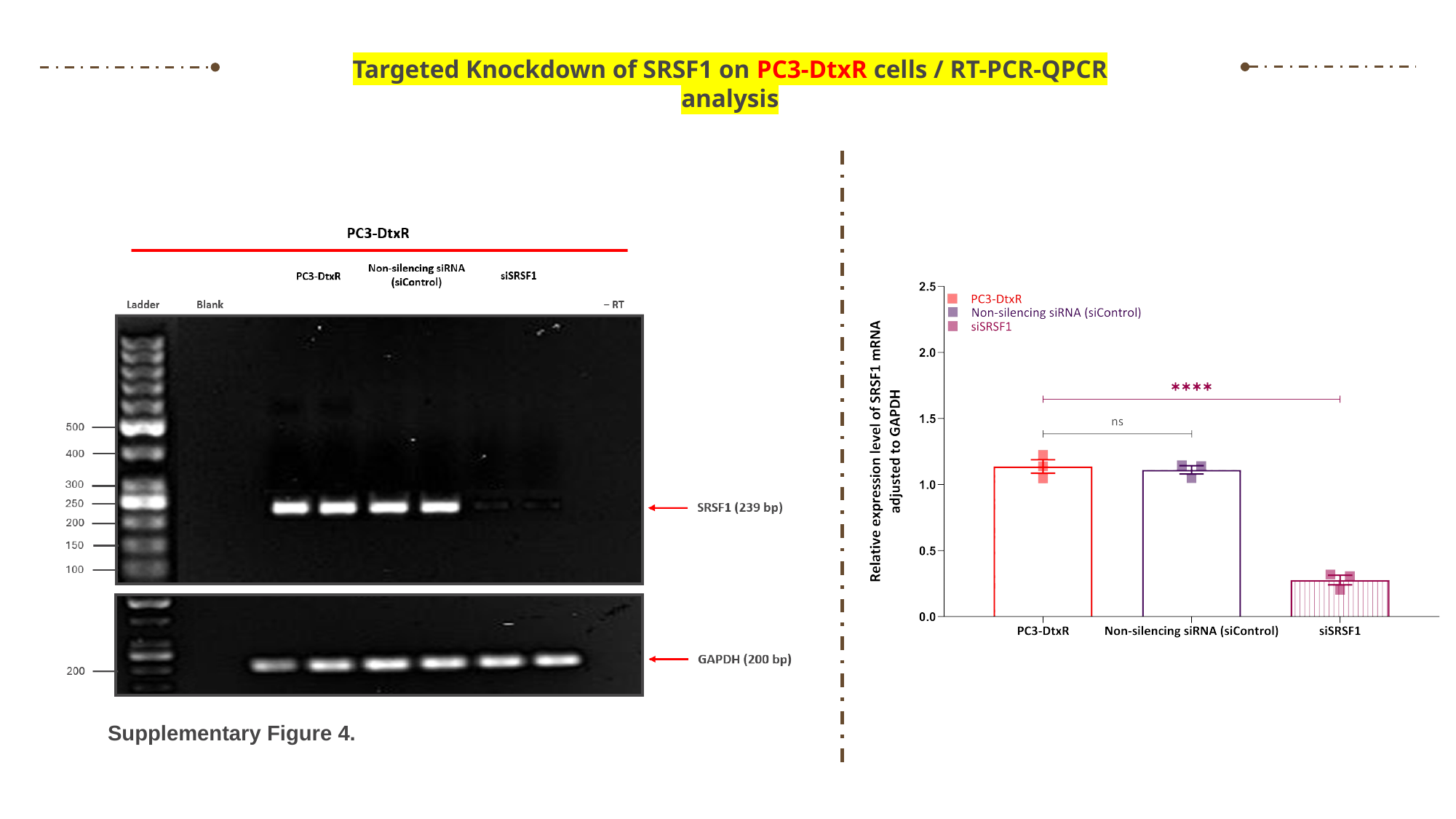

Targeted Knockdown of SRSF1 on PC3-DtxR cells / RT-PCR-QPCR analysis
Supplementary Figure 4.

### Slide 5
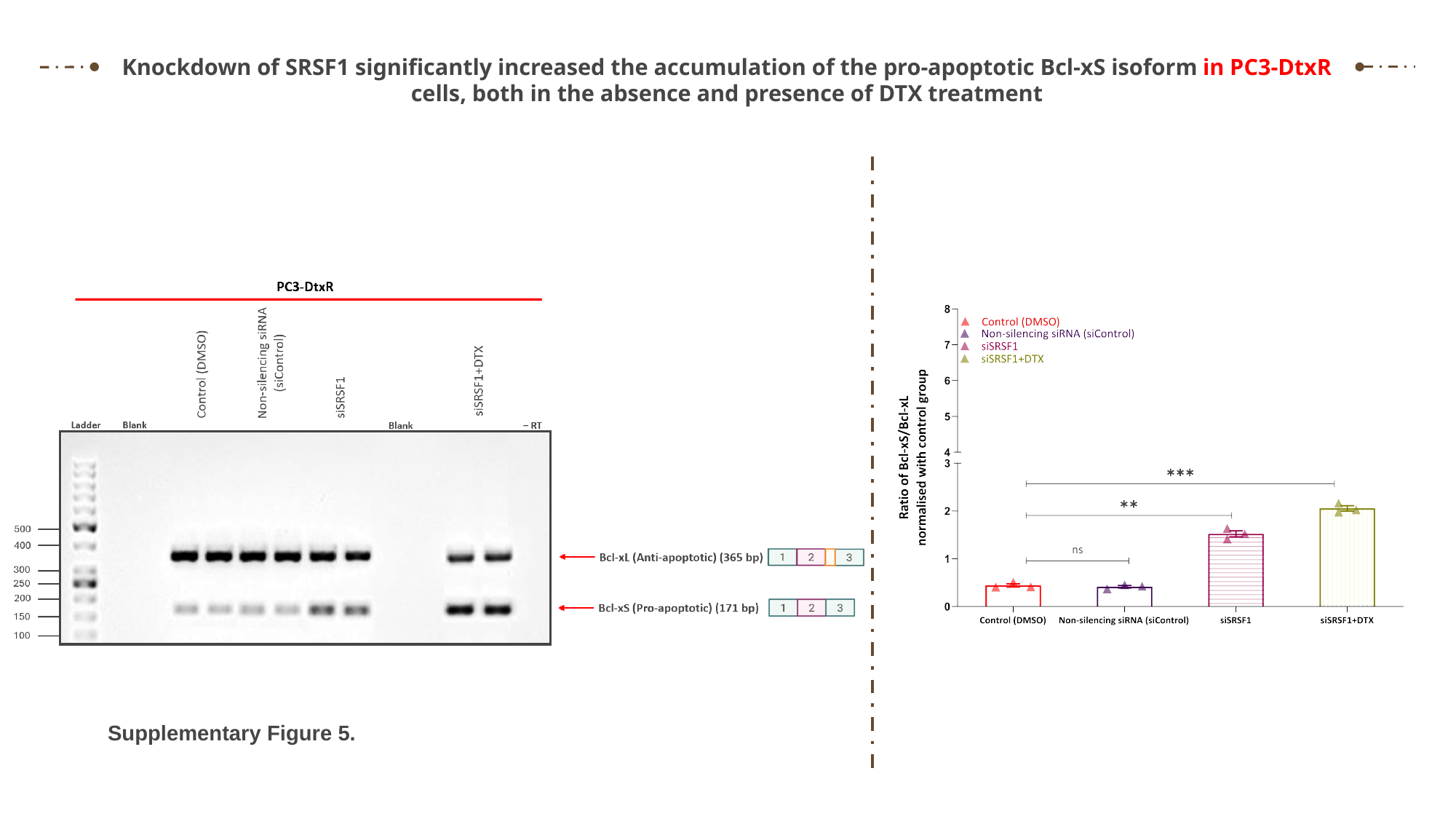

Knockdown of SRSF1 significantly increased the accumulation of the pro-apoptotic Bcl-xS isoform in PC3-DtxR cells, both in the absence and presence of DTX treatment
Supplementary Figure 5.

### Slide 6
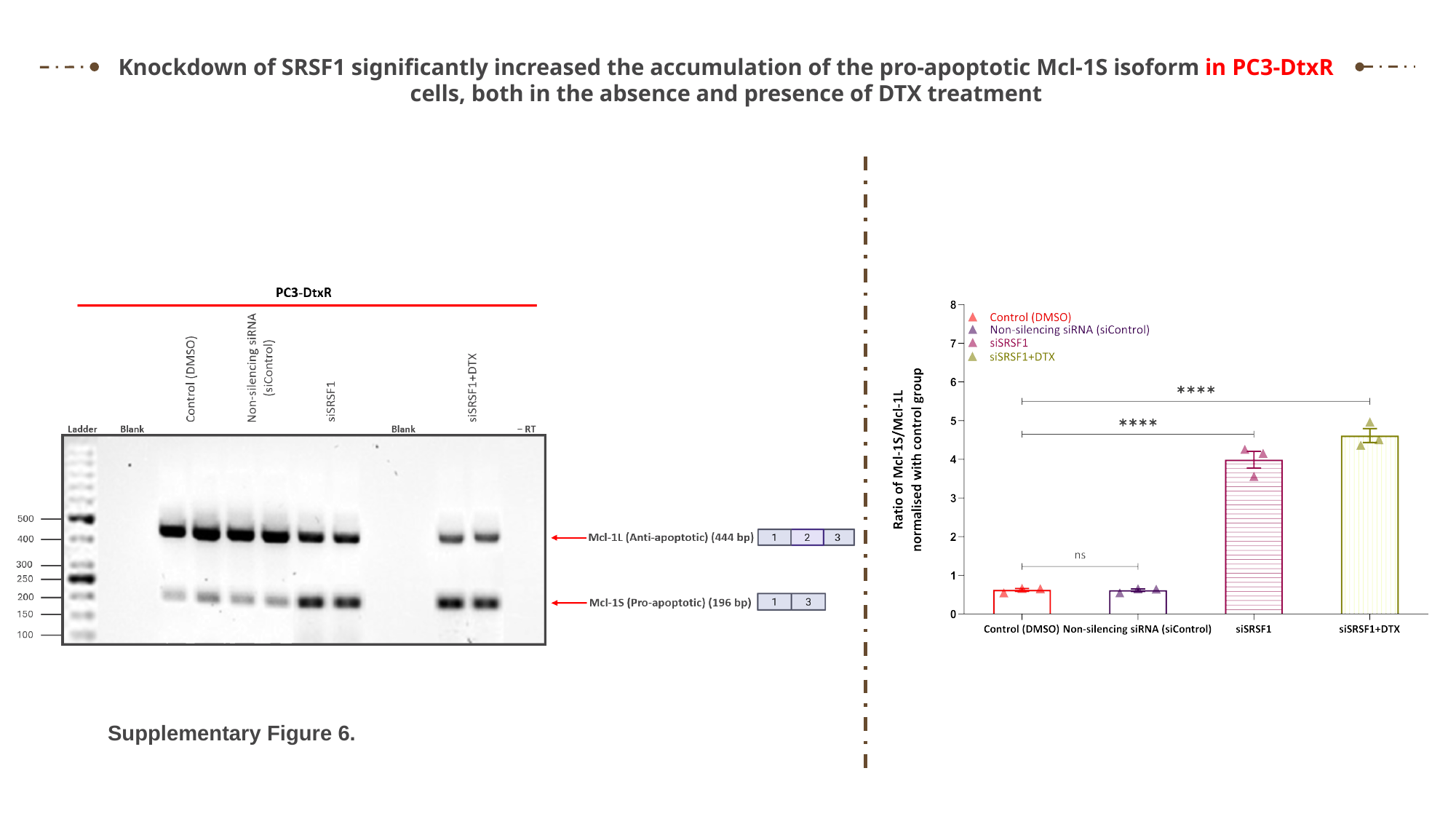

Knockdown of SRSF1 significantly increased the accumulation of the pro-apoptotic Mcl-1S isoform in PC3-DtxR cells, both in the absence and presence of DTX treatment
Supplementary Figure 6.

### Slide 7
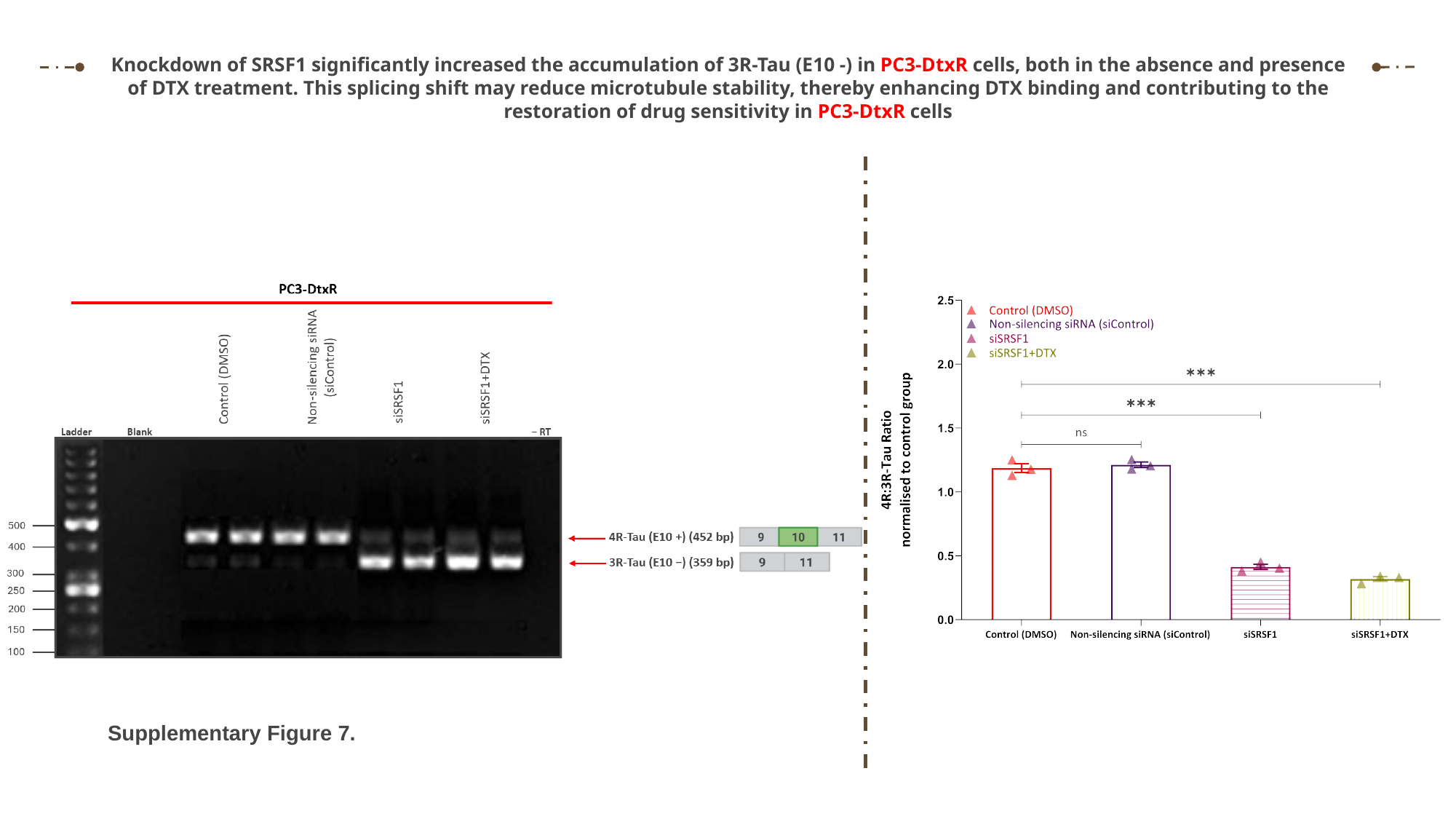

Knockdown of SRSF1 significantly increased the accumulation of 3R-Tau (E10 -) in PC3-DtxR cells, both in the absence and presence of DTX treatment. This splicing shift may reduce microtubule stability, thereby enhancing DTX binding and contributing to the restoration of drug sensitivity in PC3-DtxR cells
Supplementary Figure 7.
